# Orthogonal representational geometry in dACC underpins human hierarchical reasoning

**DOI:** 10.1101/2025.11.16.688413

**Authors:** Chuanyong Xu, Ning Mei, Wenshan Dong, Rongcheng Hu, Tom Verguts, Qi Chen

**Affiliations:** School of Psychology, Shenzhen University, Shenzhen, China; School of Psychology, Center for Studies of Psychological Application, and Guangdong Key Laboratory of Mental Health and Cognitive Science, South China Normal University, Guangzhou, China; Department of Experimental Psychology, Ghent University, Ghent, Belgium

**Author notes:** Correspondence (Q.C.).

**Keywords:** hierarchical reasoning, dACC, RNN, orthogonal representational geometry

## Abstract

Flexible adaptation requires inferring the causes of feedback and adjusting accordingly. When tasks are hierarchically structured with high-level rules governing low-level perceptual judgements, good performance requires hierarchical reasoning. Yet the computational and neural mechanisms underlying this process remain unclear. Here, building on Bayesian modelling and recurrent neural network simulation of hierarchical reasoning, we verified that such algorithm and representations are implemented in human dorsal anterior cingulate cortex (dACC). dACC encodes accumulated errors and perceptual difficulty to estimate high-level rule switch confidence. These two variables were represented along two separable dimensions, and could be approximated by Gaussian basis functions. Orthogonality between these two dimensions yielded an optimal two-dimensional representation for hierarchical reasoning. A closer-to-orthogonal representational geometry also predicted better performance. Together, these findings provide an integrated computational and neural representational basis for hierarchical reasoning.

## Introduction

Humans possess a remarkable ability to infer the sources of external feedback, and adaptively adjust their behavioural strategies to achieve cognitive flexibility ^1–3^. They must often operate in noisy and dynamic environments, in tasks with hierarchical structure, wherein high-level stimulus–response rules determine low-level perceptual decisions. In such situations, the causes of (negative) outcomes are ambiguous: One needs hierarchical reasoning to infer whether an error arises from low-level perceptual noise or instead from a high-level rule change ^2–6^. For example, when facing communication difficulties in an unfamiliar city, one may infer whether the problem arises from misperceiving speech or from differing regional dialects. This form of reasoning enables one to flexibly update behavioral strategies in response to environmental changes. Despite its importance, the computational and neural mechanisms underlying hierarchical reasoning remain unclear.

At the neural level, classical cognitive theories and functional magnetic resonance imaging (fMRI) findings suggest that the prefrontal cortex engages in hierarchical reasoning ^7–15^. In particular, the dorsal anterior cingulate cortex (dACC) is thought to play a central role in tracking errors ^12^ and estimate expected outcomes and confidence ^9,16^. Nevertheless, how exactly hierarchical reasoning is computationally implemented in the dACC remains an open question.

Bayesian modeling offers a useful computational framework for understanding hierarchical reasoning. According to Bayesian principles ^17^, humans integrate accumulated errors with current accuracy of low-level perceptual judgments to dynamically update posterior beliefs about high-level rules. This process involves accumulating confidence in the need for a rule switch, which in turn guides adaptive adjustments in behavioural strategy ^2,3,5,18,19^.

While a Bayesian model can provide an algorithmic-level account of hierarchical reasoning, neural network modelling offers a pathway to uncover its implementation ^20,21^. Because they can maintain and process information over time, recurrent neural networks (RNN) can effectively simulate human reasoning in complex cognitive tasks^8,21–23^. This is particularly evident when the RNN hidden layer exhibits basis function properties, enabling their computations to approximate Bayesian inference ^17,24,25^. We therefore adopt an RNN framework to simulate hierarchical reasoning, leveraging a representational perspective that focuses on neural patterns and their geometry ^21,26–29^.

RNNs allow studying representational patterns across task conditions, and representational geometry captures the structural relations among those patterns (e.g., distances and angles) ^30–32^. A widely observed form of representational geometry is orthogonality, which can minimize interference between task variables ^33–35^, promote knowledge generalization across contexts ^8^, and support hierarchical organization of sequential contents in working memory ^36^. Inspired by these findings, we hypothesize that an RNN trained to simulate hierarchical reasoning will develop orthogonal dimensions in its hidden layer, separating task-relevant information and forming an optimized hierarchical structure for reasoning. Furthermore, we predict that human brain regions involved in hierarchical reasoning, will rely on a similar representational mechanism.

To test these hypotheses, we employed a hierarchical reasoning task and combined Bayesian computational modeling, RNN simulation, and fMRI. We fitted a confidence-based Bayesian model that estimates rule-switch probability based on accumulated errors and perceptual difficulty. In the RNN, we observed that these variables were encoded along two dimensions of hidden layer unit activity, with patterns that approximated Gaussian basis functions. Critically, the geometry between these two dimensions was orthogonal, which can support optimal hierarchical representations and flexibly track rule-switch probability during reasoning. Guided by these model predictions, we then identified the human brain regions implementing such a mechanism. Univariate fMRI analyses showed that the dACC engaged in hierarchical reasoning about sources of feedback, jointly encoding accumulated errors and perceptual difficulty to estimate rule-switch confidence. Multivariate analyses further confirmed, within dACC, a two-dimensional hierarchical representational structure consistent with the RNN hidden layer. Furthermore, subjects’ performance during hierarchical reasoning was significantly predicted by their degree of orthogonality in their geometry. Together, these results provide direct evidence that orthogonal representational geometry supports hierarchical reasoning.

## Results

### Hierarchical reasoning based on accumulated errors and perceptual task difficulties

To characterize behavior when humans perform hierarchical reasoning, we conducted a rotation judgement task which required subjects to judge the rotation direction of two sequentially presented gratings. The rotation angle intervals were determined per subject using psychophysical methods (see Supplementary Figure 1) to obtain precisely calibrated difficulty levels. Specifically, we selected the rotation angle corresponding to probabilities of 0.1 (easy), 0.3 (middle), 0.5 (hard), 0.7 (middle), and 0.9 (easy) anti-clockwise judgements (i.e., **Figure 1b** x-axis) to serve as difficulty indicators in the main experiment. The main task was implemented in a volatile environment with two alternating stimulus-response rules. In each trial, subjects made a rule judgment (red/blue) followed by a perceptual judgment of the grating rotation (clockwise vs. anti-clockwise), conditional on the (red or blue) rule. Reward was delivered only when both rule and perceptual judgments were correct; otherwise, negative feedback was presented. During the red rule, if the test grating was rotated clockwise relative to the reference grating, subjects had to press “1” with their right index finger; if it was rotated anti-clockwise, they had to press “2” with right middle finger. For the blue rule, the key press mappings are reversed (**Figure 1a**). The two rules changed in a blocked fashion, with a minimum of 5 trials plus a sample from a geometric distribution with a mean of 4 trials. In the inference rule task (**Figure 1a** upper), no rule cue was shown, and subjects had to infer the rule based on trial-by-trial feedback. When an error occurred, subjects had to reason about the cause of this error (rule or grating rotation), and decide to either switch rules or instead optimize their perceptual judgement. As a control condition, in the instructed rule task, the currently-relevant rule was explicitly cued before each perceptual judgment (**Figure 1a** lower).

**Figure 1.**
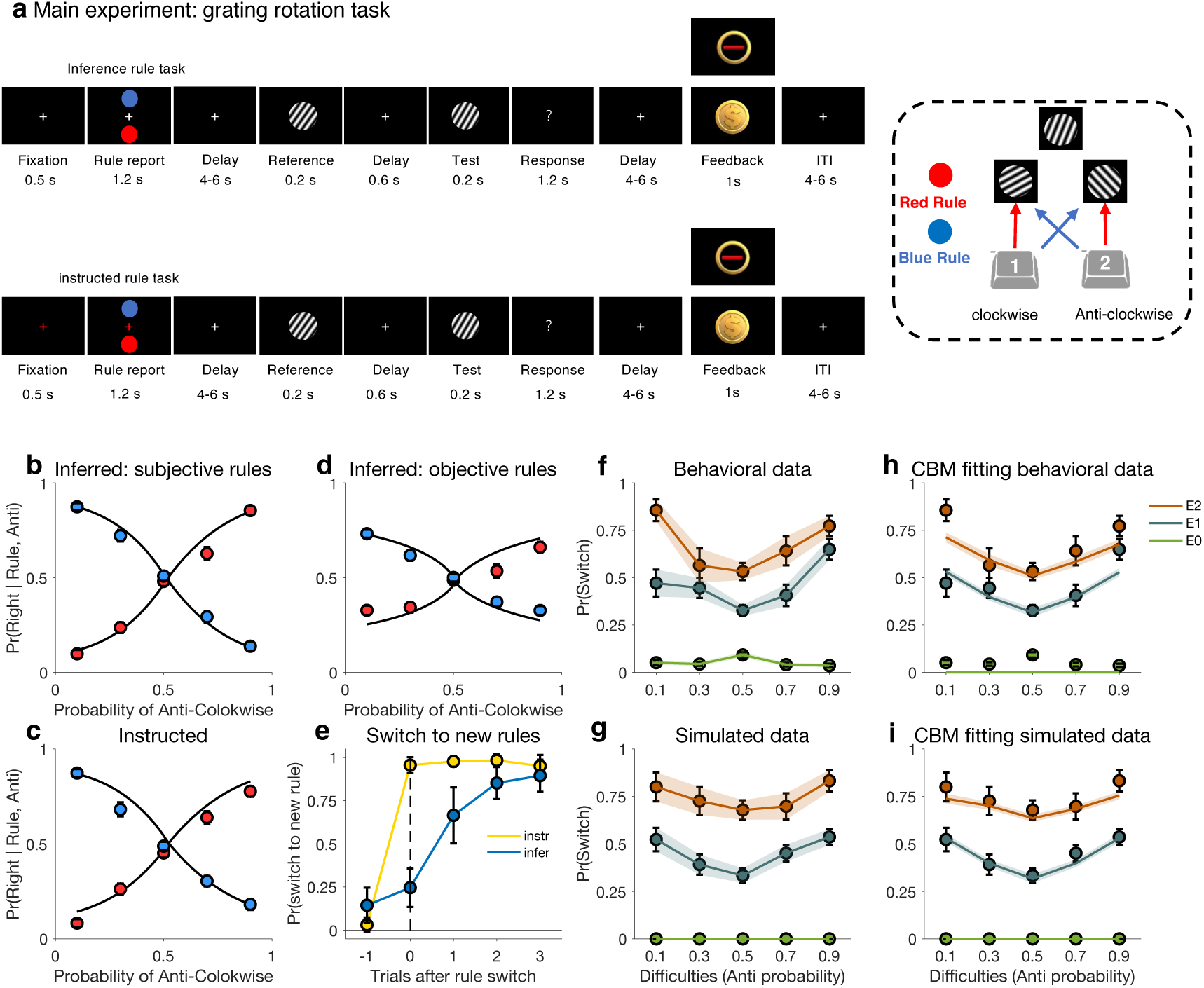
Hierarchical reasoning experimental paradigm and behavioural results. **a,** (left) Experiment paradigm for rule inference (upper) and rule instructed (lower) task. Subjects infer the rule (red/blue), then discriminate if the test grating is clockwise- or anti-clockwise rotated compared to the reference grating. If subjects receive an error (red line) at the feedback screen, they need to disambiguate its cause (rule or perceptual error). (right) Task contingencies: stimulus-action mapping under red and blue rule. **b,** Probability of a right response as a function of subjects’ reported rule (“subjective”), angle between reference and test gratings. **c,** Same as (b) for the instructed rule task. **d,** Same as (b), but for experimentally imposed rule (“objective”). **e,** Probability of switching to new rule by subjects after the objective rule changed, for the instructed (yellow) and inferred (blue) task. **f,** The probability of switching to a new rule as a function of difficulties and consecutive errors. Error bars and lines show subjects’ performance. The probabilities (anti) of 0.1 (easy), 0.3 (middle), 0.5 (hard), 0.7 (middle), and 0.9 (easy) anti-clockwise judgements serves as difficulty indicators in x-axis. E0, without error; E1, first consecutive error; E2, second consecutive error. **g,** Same as (f) for simulated data. **h,** Same as (f), but here lines show CBM model fits. **i,** same as (g), but here lines show CBM model fits. CBM, confidence based model.

To assess behavioural performance, we first measured the proportion of Right (i.e., “2”) key presses as a function of the rule and rotation angle (corresponding to probabilities of anti-clockwise judgements), or Pr(Right | Rule, Anti). In the instructed rule task, this probability increased as a function of angle with the red rule and decreased for the blue rule (**Figure 1c**). In the inference rule task, we analyzed key presses both in terms of the subjective rule reported by subjects (**Figure 1b**) and the objectively active rule (**Figure 1d**). Performance in the inference task as computed conditional on the subject’s inferred rule was close to the instructed rule task, indicating that subjects understood and responded to the hierarchical structure of the task. Performance on the inference task was of course worse when conditioned on the objective rule, because subjects could make incorrect inferences on the currently active rule.

In the instructed rule task, subjects switched their choice of rule immediately after the active rule switched (**Figure 1e**). In the inference rule task, the rule was implicit, so subjects had to infer the rule switch from the pattern of feedback. The rule switch inference could be based on accumulated errors (number of errors), and further modulated by the probabilities of anti-clockwise judgements (i.e., perceptual difficulty; **Figure 1f-i**) ^3,5^. We collapsed the 5 probabilities of anti-clockwise judgements into three difficulty levels: hard, middle, easy, and labeled them as difficulty levels 1, 2, 3, respectively, for statistical analysis. Quantitatively, mixed-effects logistic regression analysis using ‘fitglme’ in MATLAB for error trials, showed that the probability of a subject switching the rule was related to both accumulated number of errors and task difficulty, in a positive manner (*β*_errors_ = 0.90, 95% CI = [0.58, 1.22], *P* < 0.001; *β*_*difficulty*_ = 0.46, 95% CI = [0.29, 0.63], *P* < 0.001). When subjects receive rewards, no rule switch is needed; an error may instead indicate that a covert rule switch occurred, especially when there are consecutive errors, and especially when the perceptual difficulty was easy (**Figure 1f**).

### Subjects update rule switch confidence based on Bayesian principles

The confidence-based model (CBM) ^5^ was used to identify the latent variables guiding subjects’ rule switch decision. The CBM first estimates the posterior probability of a rule switch by integrating the expected accuracy of the current perceptual judgement with the outcomes of preceding trials. Building on this, the CBM summarizes the accumulated evidence for a covert rule switch into a single latent variable, *C*_switch_. After updating *C*_switch_, the model issues a binary switch decision depending on whether *C*_switch_ is larger than a threshold, θ (**Figure 1a** lower right). We included a hazard rate parameter ^18^ and a perseveration factor ^5^ to optimize the model, because these factors enabled us to account for systematic deviations from the normative Bayesian prediction about rule switch. Model fitting and simulation results demonstrated that the CBM successfully captured performance in this task (**Figure 1f-i**). Model comparison further established that the CBM fitted better than the Rescorla-Wagner model ^37^, the adaptive learning rate (ALR) model ^38^, and the Sync model ^39,40^ in our data (Supplementary Figure 2).

### Recurrent neural network simulation suggests a basis function representation for hierarchical reasoning

Bayesian inference via confidence computation is a useful principle for inferring rules and learning from ambiguous feedback. However, the complexity of such confidence computations renders them implausible as a mechanism that the human brain would use, given the brain’s limited computational and storage abilities ^25^. For this reason, we next turned to the basis function representational framework, which provides a computationally efficient approach for integrating information ^41^ and subsequent linear or nonlinear transformation to approximate intractable variables such as confidence or uncertainty in Bayesian inference ^25^. Importantly, basis functions can be efficiently implemented in recurrent neural networks ^42,43^. We further hypothesize that an RNN hidden layer can approximate basis functions, and that in this way RNNs can effectively simulate hierarchical reasoning.

Here, we trained an Elman RNN with a single, 32-unit hidden layer to predict the rule switch probability for the inference rule task. As input, the RNN received a 3-dimensional stimulus-difficulty vector and a one-hot feedback vector (**Figure 2a**). The RNN processed this sequence of inputs through its nonlinear, recurrent hidden dynamics, which enabled it to approximate a basis function network ^41^. Results showed that the rule switch probability of the RNN (lines) was close to that of subjects (error bars), *R^2^* = 0.68 (**Figure 2b**). To characterize the RNN’s representations, we extracted each hidden unit’s activation at every error level, yielding a 32 (number of units) × 3 (error level: E0/E1/E2) matrix. Activations were min-max normalized across error levels. We then identified all units whose responses peaked at a specific error level. The normalized response of all units within each error level were subsequently fitted with a Gaussian function ^44^ (**Figure 2c** left). This fitting procedure was subsequently repeated for perceptual difficulty (D1/D2/D3: hard, middle, easy level). The fitting approach was motivated by the established role of Gaussian tuning curves as a universal basis for representing and transforming probability distributions ^17,24,45^. We found that the hidden unit activation patterns were well approximated by Gaussian basis functions, with a mean *R^2^* of 0.88 across the error levels and 0.82 across the difficulty levels. This representation in turn can support confidence-based inference by combining (multiplying) them (**Figure 2c** right).

**Figure 2.**
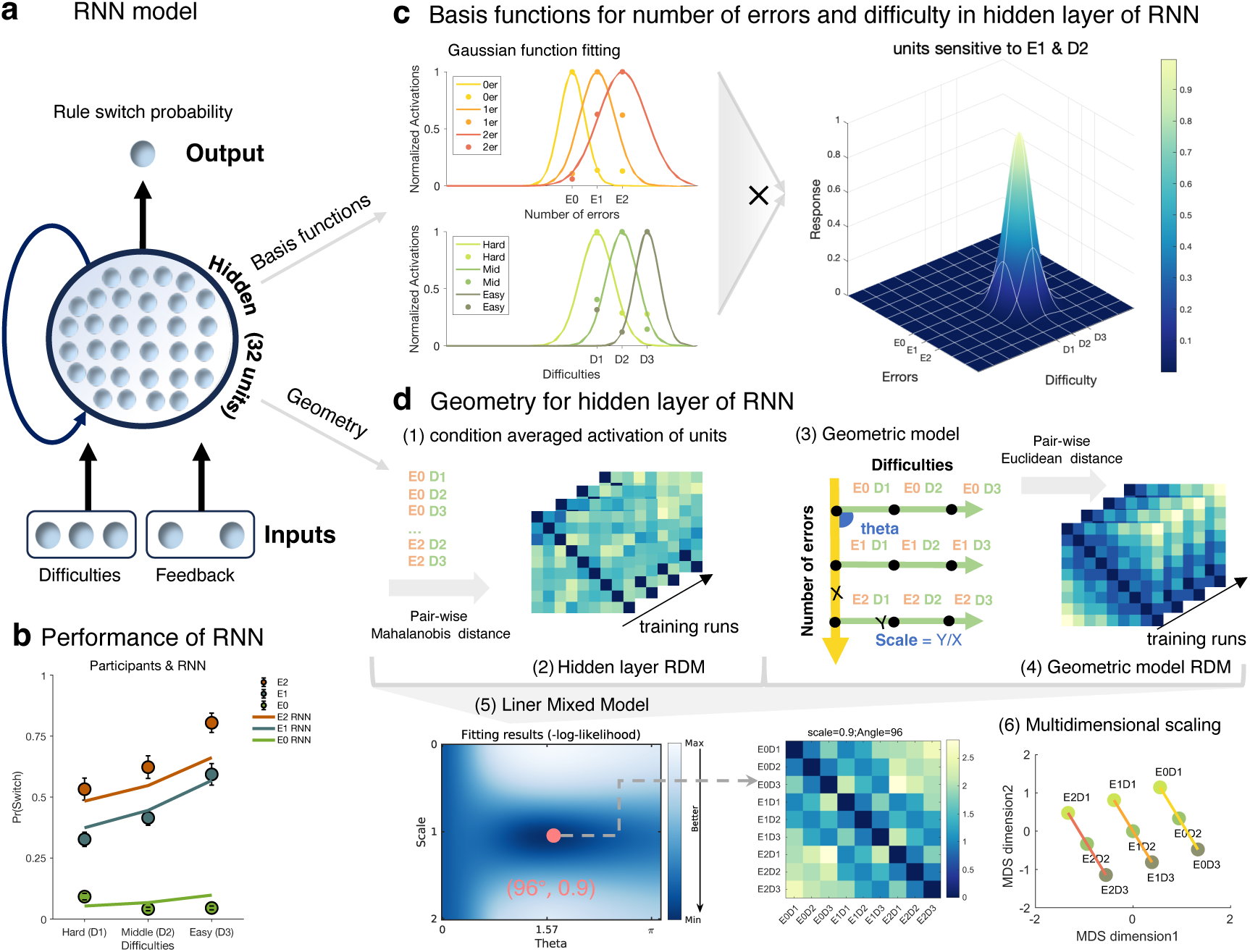
The representational geometry in the RNN hidden layer supports hierarchical reasoning. **a,** Diagram of the recurrent neural network (RNN). The RNN received perceptual difficulties (Hard: 1 0 0, Middle: 010, Easy: 0 0 1) and feedback (reward: 1 0, error: 0 1) as input for predicting the trial-by-trial rule switch probability of each subject. **b,** The RNN’s predicted rule switch probability (lines) closely matched the human data (error bars). **c,** Representational patterns for number of errors and task difficulty in the hidden layer of the RNN. We identified the units with maximal activation for each variable level. For example, some units respond maximally to 2 errors (red point on the right), moderately to 1 error (red point in the middle), and minimally to zero errors (red point on the left). The activation of these units to each error level can be fitted by a Gaussian function (red curve) *f*_*errors*_(*x*). We repeated this fitting procedure for all units. Then we calculated the product *f*_*errors*_(*x*) × *f*_*difficulty*_(*x*) to simulate the ideal global activity pattern (right). **d,** Hidden unit activations were extracted for fitting the geometric model, followed by visualization using MDS. (1) Hidden unit activations under 9 conditions were extracted to obtain the hidden layer RDM. (2) The hidden layer RDM were concatenated over training runs. (3) The geometric model for building the geometric model RDM. (4) Geometric model RDMs were concatenated. (5) (lower left) Then, linear mixed model was applied to search the best fitting (minimum −loglikelihood) theta and scale parameter of the geometric model on the hidden layer RDM. (lower right) The best fitting geometric model for the RNN hidden layer was almost orthogonal (scale=0.9, theta=96° × π/180°). (6) The multidimensional scaling for recovering the low-dimensional geometry of errors and difficulty; points reflect the 9 specific conditions. Results showed the geometric model in RNN hidden layer was similar to human subjects (see Figure 5).

To quantify the geometry underlying these representations, we next fitted the hidden layer activations with a parametric geometric model (**Figure 2d**). This geometric model defines a Euclidean two-dimensional space, which is designed to disambiguate the sources of error. An observed error can arise from either of two causes: low-level perceptual noise (captured by the difficulty dimension) or from using a high-level incorrect rule. Flexibly combining these two sets of information is important to infer the most likely cause of an outcome and compute a rule-switch probability. The geometric model is parameterized by the angle (theta) between the two axes (dimensions) and by a parameter that scales their relative units ^36^. Angle controls the separability of the two dimensions, where orthogonality (θ ≈ 90°) minimizes interference. The scale parameter controls the dimensions’ relative contributions, determining how strongly each influences the combined representation. An orthogonal geometry (θ ≈ 90°, scale ≈ 1) minimizes interference, allowing for an optimal two-dimension hierarchical structure for reasoning.

We next compared the geometric model with the RNN hidden layer via representational similarity analysis (RSA). The Euclidean distances between each pair of 9 conditions (3 error levels × 3 difficulty levels) were calculated based on the geometric model, to obtain a first representational dissimilarity matrix (gRDM) (**Figure 2d**). The Mahalanobis distance between each pair of 9 condition-averaged activations in the hidden layer in each training run were also calculated to obtain a second RDM, called the hidden layer RDM. Then the hidden layer RDMs were concatenated and regressed against the gRDM using a Linear Mixed Model. We fit a grid of linear mixed models, where the geometric model was parameterized by the inter-axis angle and scale ratio, selecting the best geometric model with the lowest negative log-likelihood (**Figure 2d**, red points). Model fits showed that the angle between the number of errors and perceptual task difficulty axes was close to a right angle (96°), and the axes were almost equally scaled (scale = 0.9) (**Figure 2d**, lower middle), consistent with near-orthogonal scaling.

As an alternative way to visualize hidden unit space, we next used multidimensional scaling (MDS) on the hidden layer unit activations. Here, the number of errors and difficulty were encoded along two orthogonal dimensions (**Figure 2d** lower right), yielding a flexible basis for rule switch probability computations during hierarchical reasoning.

### Inferring the sources of feedback engages dACC and superior prefrontal gyrus

To understand how the brain conducts hierarchical reasoning, we next tried to identify the regions engaged in feedback inference, particularly those involved in error attribution. Because the inference and instructed rule task differ only in whether the rule must be inferred, contrasting their feedback epochs isolates activity specific to feedback-based inference. Accordingly, we applied two condition-specific general linear models (GLMs) to the fMRI data to estimate activation strength during feedback inference.

After extracting beta maps from the GLM _inference_ and GLM _instructed_, a first *t*-test contrast revealed that feedback onset evoked stronger activation in dACC (peak *t*_28_ = 4.56, MNI =[-6 22 44], cluster size = 417), right insular (peak *t*_28_ = 4.93, MNI = [32 20-4], cluster size = 317), and right superior prefrontal cortices (peak *t*_28_ = 4.86, MNI =[20 26 54], cluster size = 262) during the inference task compared to the instructed task (**Figure 3a**). In the second *t*-test, we contrasted error feedback onset in the inference versus instructed task (**Figure 3b**). The dACC (peak *t*_28_ = 4.77, MNI = [-6 22 42], cluster size = 189) and right superior prefrontal cortices (peak *t*_28_ = 4.87, MNI = [20 26 52], cluster size = 181) showed statistically significant evoked responses. These results all survived at cluster-based FWE correction (voxel level *P* < 0.001) for multiple comparisons.

**Figure 3.**
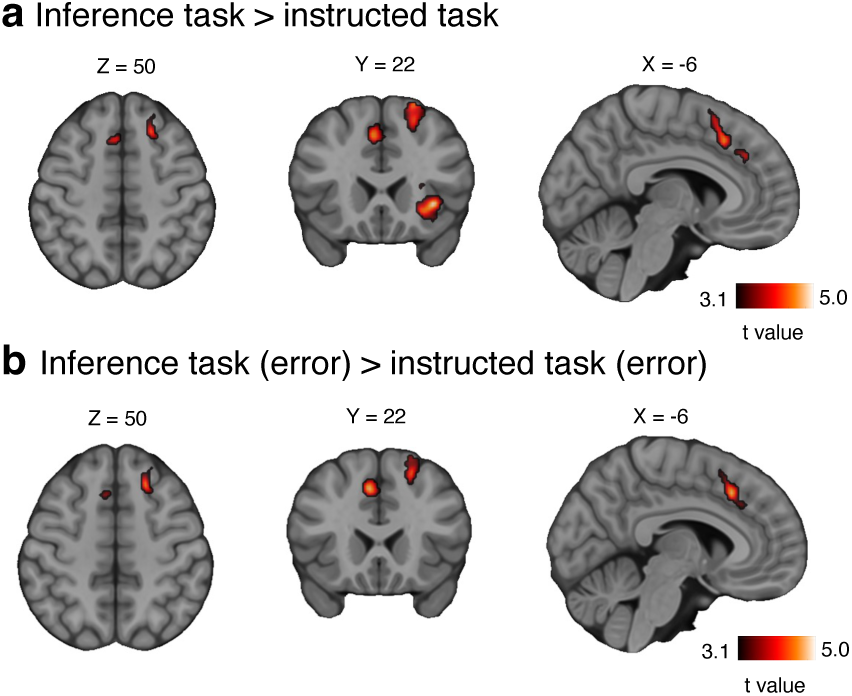
Human brain activation at feedback error inference. **a,** Main effects for task type, contrast between rule inference and instructed task. The results were first thresholded at a voxel-level of *P* < 0.001, followed by a cluster correction procedure with a cluster level threshold of *P* < 0.05 (FWE). Only clusters size larger than 262 voxels were retained. **b,** Stronger activity in the inference task than in the instructed rule task, when subjects receive error feedback. Cluster-based FWE correction was also applied; only clusters larger than 181 voxels retained. The cross-section at MNI X = −6, Y = 22, Z = 50.

### Human dACC encodes accumulated errors and perceptual difficulties for rule switch confidence

The CBM posits that evidence for a rule switch during hierarchical reasoning is encoded in terms of confidence, which depends on the number of consecutive errors and task difficulty. To localize the brain regions engaged in these variables during the inference rule task, three parametric modulation GLMs were constructed for number of errors (GLM-PM1), perceptual difficulty (GLM-PM2) and confidence (GLM-PM3) separately (**Figure 4a**). Modeling them in separate GLM-PMs avoids multicollinearity, stabilizes estimates, and circumvents order-dependent orthogonalization settings. This design ensured the estimated total effect of each variable on the BOLD signal was not confounded by its shared variance with the others. For all three GLM-PMs, the BOLD signal was regressed on the feedback screen. A one-sample *t*-test against 0 revealed that a positive modulation effect of errors in dACC (GLM-PM1; peak *t*_28_ = 6.80, MNI = [-4 18 42], cluster size = 1251), right middle prefrontal cortices, left thalamus, and right thalamus, survived cluster-based FWE multi comparations correction (voxel level *P* < 0.001). Only the negative modulation effect of difficulty survived in dACC (GLM-PM2; peak *t*_28_ = −4.74, MNI = [8 20 40], cluster size = 379), and right middle prefrontal cortices. The rule switch confidence parameter extracted from the CBM positively modulated the response in dACC (GLM-PM3; peak *t*_28_ = 5.38, MNI = [-6 18 44], cluster size = 227). More details and negative modulation effects for errors and confidence are reported in Supplementary Table 2. The peak cluster encoding errors, difficulty, and confidence mostly overlapped (**Figure 4b**). This suggests dACC is a hub for integrating the accumulation of errors and task difficulty for the purpose of computing rule switch confidence.

**Figure 4.**
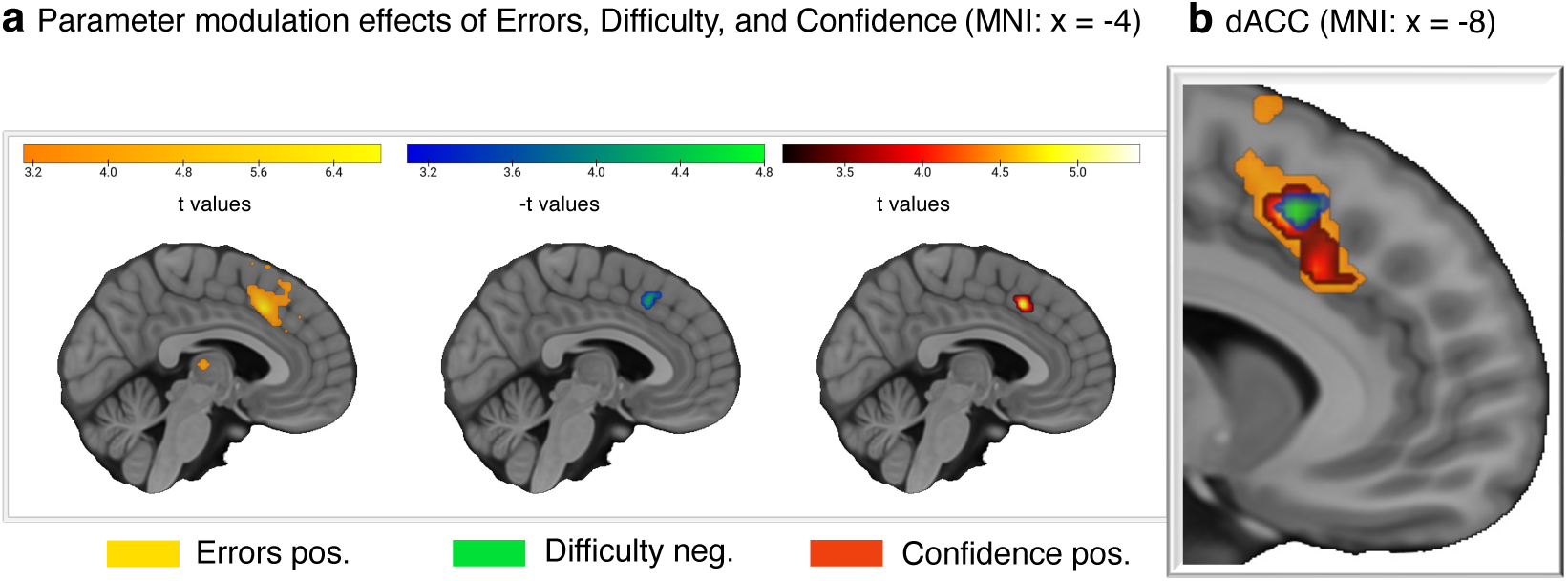
dACC activation is modulated by key variables for rule switch confidence. **a,** Parametric univariate modulated activation from number of errors (yellow), perceptual difficulty (green), and rule switch confidence (red). The peak coordinates occurred in dACC. **b,** A zoom of dACC showing the modulation effects of panel a. The cluster of errors, difficulty, and confidence effect overlapped at the superior part of dACC.

The functional and anatomical organization of the dACC exhibits a gradient-like structure that supports information integration, decision-making and cognitive control ^10,33,46–48^. Consistently, in our data the conjunctive maps showed a gradient of increasing encoding strength from the inferior to the superior part in dACC. Errors, difficulty and confidence were most strongly encoded in superior dACC (**Figure 4b** right). The overlapping superior part in dACC for errors and difficulty encoding may support the idea of two-dimensional organization ^36^ for confidence computation during hierarchical reasoning.

### dACC exhibits representations similar to the RNN

Motivated by the representations in the hidden layer of the RNN, we tested whether similar representations arise in the human brain. While the GLM univariate analysis points towards overlap of number of errors and perceptual difficulty in dACC, they don’t specify how errors and difficulty are coded ^49^. To address this, we next characterized the activity pattern of errors and difficulty in dACC, using multivariate pattern analysis (MVPA) ^50^. Because we were interested how (the representations of) errors and difficulties were integrated to support rule switch inference, all of the subsequent analyses focused on the subregion of dACC that encoded confidence (whole confidence dACC) (red region in **Figure 4b**).

MVPA was implemented using linear support vector machine (SVM). The whole confidence-related dACC could classify number of errors with higher Area Under Receiver Operating Characteristic Curve (ROC-AUC) score than random. Moreover, the superior dACC subregion identified in the encoding analysis (**Figure 4b**) can significantly classify both error count and perceptual task difficulty (**Figure 5a** middle; multiple comparison corrected at *P* < 0.05 with voxel-level FWE).

**Figure 5.**
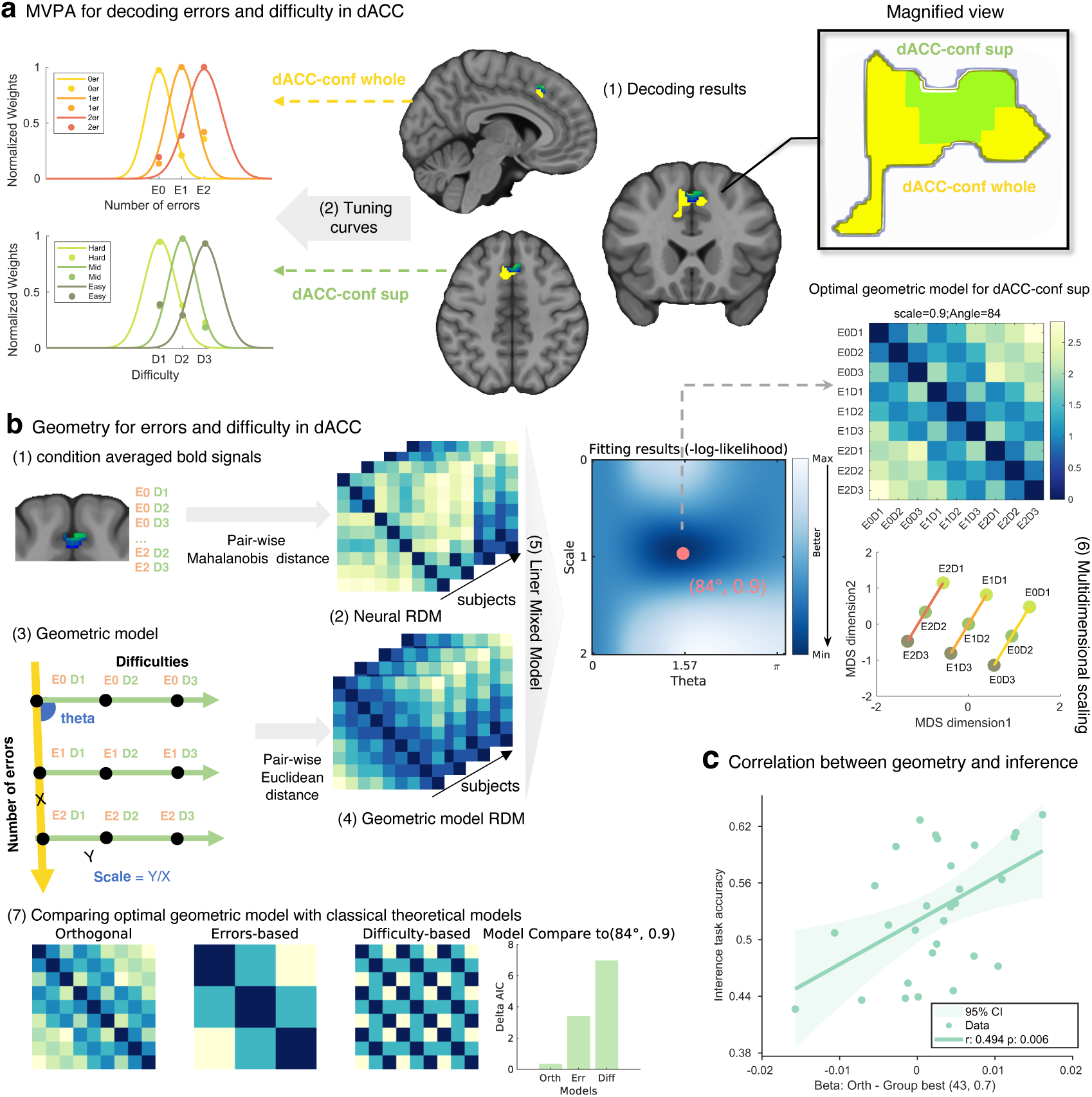
Neural representational geometry supporting the two-dimensional hierarchical representations for rule switch inference. **a,** (middle) Multivariate pattern analysis (MVPA) showed that number of errors and level of difficulties can be decoded in the part of dACC that encoded confidence. All this area can decode errors (yellow), but only the superior subregion could decode both errors and difficulty (green). (left) Decoding weights for each level of errors and each level of difficulty were extracted to plot the tuning curves, and fitted with Gaussian functions. (right) Zoom into the decoding areas. **b,** (1) BOLD signals under 9 conditions in the superior part of dACC were extracted, and the Mahalanobis distance between each pair of conditions were calculated to construct the neural representational dissimilarity matrices (RDM). (2) The neural RDM of all subjects were concatenated. (3) Geometric model for characterizing the structure of errors and difficulty, with 2 axes and 9 points by crossing errors and perceptual difficulties. The angle and length ratio between the errors and difficulty axis is determined by the angle (theta) and the scale (Y/X), respectively. The Euclidean distance between each pair of conditions were calculated to construct the geometric model RDM. (4) Geometric model RDM of all subjects were concatenated. (5) Then, a linear mixed model was applied to search the best fitting (minimum −loglikelihood) theta and scale parameter of the geometric model on neural RDM. (middle right) The best geometric model for dACC in superior part was almost orthogonal (scale=0.9, theta=84° × π/180°), which may be a necessary condition for simultaneously decoding errors and difficulty. (6) The multidimensional scaling for recovering the low-dimensional geometry of errors and difficulty in dACC. Individual points reflect the (3*3) conditions. (7) Empirical model comparison also points to an orthogonal geometry in the superior subregion of dACC. **c,** In the whole confidence-modulated dACC area, subjects with a more orthogonal representational geometry, performed the inference task better.

After training a linear SVM decoder that can discriminate the three error levels (E0/E1/E2) and the three task difficulty levels (D1/D2/D3), we next extracted searchlight averaged classification weights for each level, and projected it into the central voxel (of each searchlight sphere with 10mm radius). In line with the representational analysis in the RNN, for each variable, voxels with the highest weight for a given level were grouped; their weights were min-max normalized (across levels), averaged within a group, and fit with Gaussian functions ^44^. The resulting curves showed level-specific responses in the dACC, with robust Gaussian fits for both the number of errors (mean *R^2^* = 0.81) and perceptual difficulty (mean *R^2^* = 0.73) (**Figure 5a** left). Similar to the RNN hidden layer units (**Figure 2c** left), activation patterns within the corresponding voxel groups were approximately Gaussian-tuned to their preferred level (peak point), yielding two sets of basis functions.

### Orthogonal representational geometry supports optimal hierarchical representations

To quantitatively characterize the geometry supporting the optimal hierarchical representation of errors and difficulty in superior dACC (green region in **Figure 5a**, right), we applied the same procedure for identifying the RNN hidden layer geometry, but now to the fMRI data in superior dACC. The Mahalanobis distance between each pair of 9 condition-averaged activations (during feedback) in superior dACC was also calculated (neural RDM). Fitting results indicated that the geometry in superior dACC is best described by a near-orthogonal arrangement (angle = 84°, scale value = 0.9), placing errors and difficulty on approximately orthogonal axes of a 2-dimensional structure (**Figure 5b** upper right). For visualizing the near-orthogonal geometry of dACC in low-dimensional space, we used MDS. The results indicated that errors and difficulty co-existed in an orthogonal neural representational space of dACC (**Figure 5b** lower right). Such a pattern was only observed in the superior part of the dACC (**Figure 5a** right, green color), which was capable of linearly decoding both variables in the MVPA performed above.

To benchmark against canonical theoretical schemes, we compared Akaike Information Criterion (AIC) values between the best fitting geometric model and alternative theoretical templates (smaller delta AIC implies greater similarity). The orthogonal template yielded the smallest delta AIC =0.33, outperforming an errors-based model (delta AIC = 3.40) and a difficulty-based model (delta AIC = 6.96), corroborating an orthogonal geometry in dACC. (**Figure 5b** lower left).

We next tried to link the neural representational geometry to each subject’s behaviour. For this purpose, we calculated the Pearson correlation coefficients between each subject’s inference task accuracy and their beta coefficient differences (theoretical orthogonal model template minus the best fitted model) in the whole confidence-related region of dACC. Beta coefficient differences with larger positive values indicate stronger alignment with the theoretical orthogonal model. In the whole confidence-related dACC region, this relationship was positive (r = 0.494, *t*_27_ = 2.95 *P* = 0.006). In other words, during feedback inference, subjects with more orthogonal dACC representations performed better in the inference task (**Figure 5c**).

## Discussion

The present study advances understanding of the computational principles and neural representations of hierarchical reasoning. At the algorithmic level, individuals update confidence in a rule switch according to Bayesian principles. This confidence depends on both accumulated errors and accuracy of low-level perceptual judgements at various levels of task difficulty. At the representational level, RNN simulation and fMRI results converge on a common mechanism. Both the RNN’s hidden layer units and the human dACC rely on a two-dimensional representation. Here, errors and difficulty are encoded along two dimensions that can be approximated by Gaussian basis functions. Critically, the orthogonality of these two dimensions supports an optimal hierarchical representation and predicts individual performance.

Consistent with prior work, dACC monitors errors and tracks behavioral outcomes over time ^12,14^. We extend this known role of dACC to hierarchical reasoning ^3,5^, and demonstrate how such variables are integrated with low-level perceptual difficulty ^5,51^. Our findings provide direct evidence that the human dACC guides behavioural strategy adjustments by estimating subjective confidence based on a Bayesian model ^9,52,53^. Our fMRI conjunctive analysis further reveals that dACC integrates evidence across several variables in an overlapping subregion, where it is transformed into confidence according to Bayesian principles ^9,54,55^, supporting hierarchical reasoning. Such a mechanism avoids the need for multiple specialized circuits, highlighting an efficient solution that recycles neural circuitry ^56^.

We further uncover the representational patterns of hierarchical reasoning in both RNN hidden layers and human dACC. According to the Bayesian brain hypothesis ^9,17,24,41^, the nervous system represents uncertainty of information probabilistically and updates it with new evidence as it comes in. Subjective confidence is essentially such a Bayesian probability estimate ^9,54,55^. Earlier work already decomposed computationally intractable Bayesian probabilities using (linear) combinations of performance-related variables ^57^. Here, we applied this approach to different performance-relevant variables, namely number of errors and task difficulty. An important challenge for future work will be how an agent learns which variables exactly are relevant for confidence computation in different tasks ^58^.

In the current paper, combinatorial representation through basis functions plays a key role. With enough basis functions, any function can be approximated and downstream tasks approximated as linearly solvable problems ^25^. Basis function representations have been widely observed across human brain regions ^59^ and cognitive processes, including visual perception ^60^, working memory ^42^, and mathematical cognition ^43^, and recently in social decision-making ^61^. We identify Gaussian-like basis function patterns in both RNN hidden layers and human dACC, supporting the Bayesian brain hypothesis in hierarchical reasoning. This may also explain why our single hidden layer RNN suffices for Bayesian confidence-based reasoning, without separate dedicated hidden layers for confidence and performance monitoring ^62,63^.

Recent work has emphasized a geometric perspective on neural representations ^27,33,61,64–67^, investigating which geometry is suitable for human and artificial computation ^68^. Specifically, such work focused on the use of *near*-orthogonal representations, which can provide an optimal compromise between compositional representations (for efficient generalization) and conjunctive ones (for flexible task implementation). We observed such representations in dACC, although our focus was more on orthogonality than on near-deviations from it (conjunctivity). This finding points to a specific representational account for the dACC’s central role in complex cognition ^69,70^, complementing prior anatomical and functional gradients perspectives^46–48,71^. We observed that accumulated errors and perceptual difficulty are encoded along two dimensions and flexibly combined ^27,33,66,67^. Orthogonal relations between dimensions enables an optimal hierarchical structure ^36^ to make rule switch probability estimates. This orthogonal geometry in dACC may be a general mechanism supporting reasoning, cognitive control, and error monitoring ^4,5,10,14^. It also provides a potential basis for recent meta-learning theories ^47,48^ integrating dACC’s multiple functions.

Taken together, these results clarify how the brain implements hierarchical reasoning. By combining computational modelling with representational geometrical analysis, we show that a hierarchical structure equipped with orthogonal dimensions can support efficient, near optimal inference. More broadly, this framework provides potential insights for reasoning and other cognitive process that require hierarchical Bayesian reasoning for optimal performance.

## Methods

### Subjects

Thirty-one Chinese participants with no contraindications for magnetic resonance imaging (MRI) were recruited. After excluding two subjects with excessive head motion during MRI scanning (framewise displacement > 1.5 mm/degrees), data of twenty-nine (16 males; mean age 21.6, range 18-25 years) subjects were included in subsequent analysis. Subjects with no history of neurological or psychiatric disorders were recruited. All subjects had normal or corrected to normal vision. This research complied with ethical regulations and received approval form the Departmental Ethical Committee of Shenzhen University. All subjects gave written informed consent before participating in the experiments.

### Task

#### Psychophysical task for perceptual threshold setting

Before the main experiment, subjects completed the psychophysical task (Supplementary Figure 1) for measuring their subjective perceptual threshold. The subjects completed 300 trials, where in each trial they had to compare the rotation direction between two gratings (test grating relative to reference grating). They pressed the “B” key for clockwise and the “H” key for anti-clockwise. After each judgement, a coin or red line image was presented as feedback. To avoid subjects comparing only with a fixed angle instead of comparing the two grating, and to ensure a balanced orientation of the gratings, the reference grating angles were set at 65 ± 2° (pro-orientation) or 115 ± 2° (anti-orientation). The maximum angular difference between reference and test grating was 18°, with 10 evenly spaced points selected, with a step size of 4. To control for unfamiliarity or response bias, subjects start the formal task only after obtaining 90% accuracy in the practice trials. The logistic regression function in ‘fitPsyche’ toolbox ^72^ was used for fitting the probability of subjects judging anti-clockwise at each grating angle difference. The angle differences corresponding to probabilities of 0.1, 0.3, 0.5, 0.7, and 0.9 anti-clockwise judgements were then selected as the perceptual judgement difficulty indicators in the main experiment. From these, three levels of difficulty (hard: 0.5, middle: 0.3 and 0.7, easy: 0.1 and 0.9) were determined.

#### Main experiment

Each subject completed the main experiment under MRI scanning. The main experiment included two parts: the rule inference task (uncued rule, **Figure 1a**) and the rule instructed task (rule cued by blue or red color of fixation). Their order was balanced across participants. To avoid expectancy effects and make full use of the original fMRI signal, we set a jittered time interval from 4 to 6 seconds. Each task consisted of 22 blocks with 21 rule switches, and each block contains 5-10 trials. Within each block, gratings of varying difficulty appeared randomly. Each task has 150 trials, and subjects must complete a total of 300 trials. Before each task, participants are provided with the following instructions: (1) Task structure. There are two rules, red and blue. On each trial, subjects must first indicate which rule they think is currently relevant (rule judgement), followed by a grating rotation direction judgement based on the rule (perceptual judgement). (2) Rule response. If the chosen rule appears above the fixation cross, participants press the “2” key; if it appears below the fixation cross, they press the “1” key (the position of the red and blue rules is randomly balanced within subject). (3) Rule-perceptual response contingency. For the red rule, if the second (test) grating is rotated clockwise relative to the first (reference) grating, subjects press “1” with their right index finger; if it is rotated anticlockwise, they press “2” with right middle finger. For the blue rule, the key press for perceptual responses is reversed. (4) Subjects must respond quickly when the response screen appears. Only when both the rule and perceptual response are correct will the participant receive a coin reward (on the screen), otherwise a red line will be shown. (5) The rule instructed task informs subjects of the rule on the first screen of each trial; the rule inference task does not explicitly prompt the rule, and participants must infer the rule based on feedback. (6) The higher the accuracy, the higher the reward participants will receive.

### Confidence-based Bayesian model

We fit a hierarchical Bayesian model (**Figure 1a**) that jointly explains perceptual decisions, confidence, and confidence-guided rule switching. Given feedback history, expected accuracy of rotation judgements, and the subjective hazard rate λ, a rule change is selected when the posterior for the new rule exceeds that for the current rule (ref. ^5^; Supplementary Material).

At the perceptual level, optimal evidence integration for two alternatives is naturally expressed as the log-likelihood ratio (LLR) of the sensory evidence. At the rule level, optimal selection is captured by a rule-switching decision variable that accumulates across trials the posterior evidence for “switch” versus “stay”, weighting feedback by the expected perceptual accuracy from the ideal observer. We formalize this with a hierarchical latent variable, the switching confidence *C*_switch_, which is updated trial-by-trial. Following each error trial, *C*_switch_ increases by an amount that depends on the accumulated number of errors and expected perceptual accuracy on the preceding trial. The switch probability is given by

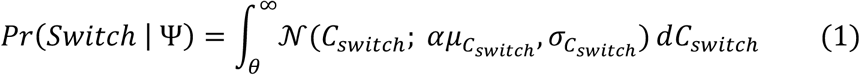

where Ψ denotes the task history, θ = 1 is the fixed threshold, μ*_C_*_switch_ is the ideal observer estimate (augmented by the perseverance factor *α* and the subjective hazard rate λ), and σ*_C_*_switch_ captures decision noise. Thus, the model has three free parameters, {σ*_C_*_switch_, *α*, λ}; a rule switch occurs when *C*_switch_ ≥ θ. The resulting rule-switch confidence is used in subsequent analysis.

### MRI sequence

Subjects were scanned with a Siemens Prisma 3T MRI system in Shenzhen University, using a 64-channel head coil. Functional blood-oxygen-level-dependent (BOLD) data were acquired using multiband gradient-echo EPI (6 acceleration factors, interleaved acquisition) with 66 axial slices covering the whole brain (phase encoding direction, anterior to posterior; Band width, 2582 Hz/Px; TR, 850ms; TE, 35ms; flip angle, 56 degrees; field of view, 210mm x 210mm; slice thickness, 2.4mm). For each task (inference and instructed rule), subjects complete three runs; each run comprised 1148 volumes. A high-resolution T1-weighted anatomical image was also acquired (TR, 2530ms; TE, 2.36ms; flip angle, 7 degrees; field of view, 256mm x 256mm; slice thickness, 1mm).

### fMRI preprocessing

The functional data of each task run was preprocessed with SPM12. Preprocessing steps for each run were slice-time correction, realigning all functional images to the first image, co-registration of the functional images to the anatomical image, normalization to standard Montreal Neurological Institute (MNI) template and resampled to 2mm isotropic resolution. Spatial smoothing was performed by using a Gaussian kernel with full-width at half-maximum (FWHM) of 6mm for general linear model (GLM) analysis in SPM, but not for multivariate pattern analysis and representational geometry analysis. MRIcroGL ^73^ was used for visualization of results.

### fMRI GLM univariate analysis

To identify brain regions that correspond to feedback and error inference, we applied the whole-brain GLM analysis. Two condition regressors were inputted in the GLM: *Y* = *X*_*errors*_*β*_1_ + *X*_*reward*_*β*_2_ + ε for rule inference and rule instructed task separately (GLM _inference_, GLM _instructed_), where *Y* represents the BOLD signal time series of a single voxel, X is the design matrix for events (a 1-s boxcar aligned to the feedback onset), β is a vector of model parameter, and ε represents the residuals. In addition to the task related feedback regressors, missing trials and nuisance regressors (6 head motion variables) were also included as covariates. Regressors were modeled separately for each run, and convolved with a canonical hemodynamic response function. Run-specific constants were included to account for between-run effects from difference of mean activation and scanner drifts. A high-pass filter (128s cut-off) was applied to remove low-frequency drifts. After the GLM model was specified and estimated, the contrast maps were extracted in a first-level analysis by applying one-sample *t*-test against 0 for each task regressor for each subject. In a second-level group analysis, the main effect contrast of task type (inference - instructed) and the simple effect of error responses (error in the inference task - error in the instructed task) were built. Statistical maps were thresholded at voxel-level of *P* < 0.001, followed by cluster-level FWE correction at *P* < 0.05.

To further identify which region’s (feedback-evoked) activity was modulated by the key variables of feedback inference, and how the rule switch confidence was encoded, we specified three parametric modulation GLM-PMs. Trial-wise number of errors (0 / 1 / 2), perceptual difficulties (1 / 2 / 3), and confidence were z-scored and entered as parametric modulators of the feedback regressors. For each modulator, first-level contrasts were estimated and submitted to second-level one-sample *t*-tests against zero.

### Representational analysis using multivariate pattern analysis (MVPA)

For multivariate decoding, unsmoothed preprocessed BOLD time series were further denoised per run by detrending, high pass filtering (128-s cutoff), z-scoring, and outlier detection (> 3 standard deviations). To avoid HRF-model assumptions, trial-wise patterns or features were defined by averaging the five volumes immediately after feedback offset, yielding one pattern per trial. Decoding performance was evaluated on the test set using ROC-AUC.

We decoded pattern differences across accumulated error levels (0 / 1 / 2) and difficulty levels (1 / 2 / 3 = hard/middle/easy) within 10-mm-radius spheres (searchlight based) constrained to the confidence-encoding mask (**Figure 4a**, right panel). Linear SVM classifiers (implemented with ‘Scikit-learn’ ^74^ and ‘Nilearn’; https://nilearn.github.io) were trained separately for error and difficulty. SVM with L1-regulation was used to reduce the probability of over-fitting, which performs well on fMRI data ^75,76^. To address the imbalance in trial numbers across error types, we up-sampled each type to match the error type with the highest trial count, using ‘Imblearn’ prior to 5-fold cross-validation, with the procedure repeated 100 times. Class probability calibration was performed using ‘CalibratedClassifierCV’.

For each searchlight, we computed the decoding precision as the mean difference between the observed decoding score and the chance level score, and assigned this value to the searchlight’s center voxel. The resulting precision maps were smoothed with sigma = 6 using FMRIB’s ‘fslmaths’. We then tested whether the precision score exceeded zero using FMRIB’s ‘randomise’ with 5000 permutations, applying voxel-wise, one-tailed inference and Bonferroni correction for multiple comparisons (*P* value < 0.05) (**Figure 5a** middle).

### Gaussian functions fitting for basis function representations

For characterizing the coding pattern of clustered voxels for different type of errors and difficulties ^32^, we extracted MVPA decoding weights for each level (errors: 0/1/2; difficulty: 1/2/3) from voxels showing significant decoding precision. For each variable, voxels were assigned to the level at which their weight was maximal (winner-take-all). Within each level-specific voxel group, weights were min-max normalized as x’ =(x-min(x)) / (max(x)-min(x)), averaged across voxels and subjects, and then fit with a Gaussian basis function to obtain tuning curves for error and difficulty ^44,77^ (**Figure 5a** left).

### Representational geometry analysis

#### Neural RDM construction

To obtain a low-dimensional visualization of the neural state space – and the representational structure defined by errors and difficulty – we computed representational dissimilarity matrices (RDMs) from condition averaged BOLD signals and applied multi-dimensional scaling (MDS). Mahalanobis distances were calculated between N condition means ^78^. Given our hypothesis that feedback inference depends primarily on error count and is modulated by perceived difficulty, we formed 9 conditions by crossing the three error levels and three difficulty levels in order: 0error-hard (E0D1), 0error-middle (E0D2), 0error-easy (E0D3); 1error-hard (E1D1), 1error-middle (E1D2), 1error-easy (E1D3); 2error-hard (E2D1), 2error-middle (E2D2), 2error-easy (E2D3).

#### Geometric model and MDS

Inspired by a 2-dimensional hierarchical representation model ^36^, and tailored to the hierarchical representation required by the present task, we defined a parametric 2-dimensional geometric model with errors and difficulty as the two axes. The geometric model is parameterized by theta (the inter-axis angle) and scale (the relative units/length ratio between the axes). For each (theta, scale) pair, we computed model distances between condition pairs as Euclidean distances in the plane, yielding a geometric model RDM. We generated a family of candidate geometries by systematically sampling parameters: theta from 0 to π radians in increments of π/180, and scale from 0 to 2 in steps of 0.1. Each model RDM was then included as fixed-effect predictor, to fit the neural RDM (or the hidden layer RDM in representational geometry analysis for RNN) at the group level using a linear mixed model (LMM). The optimal geometry was identified as the one with the minimum negative log-likelihood (**Figure 5b** middle). Pairwise distances of RDM were then embedded into a low-dimensional space using MATLAB’s ‘mdscale’, providing a summary visualization of the brain’s representational structure for these variables (**Figure 5b** right).

#### Classical theoretical model comparison

We further compared the best-fitting geometric model against three alternatives: an errors-based (1-D) model, a difficulty-based (1-D) model, and a geometric model with two perfectly orthogonal dimensions (2-D). AIC values from the LMM fits were used to compute delta-AIC relative to the optimal geometric model; smaller delta-AIC indicates greater similarity. This analysis evaluated which hypothesis best accounts for dACC neural data (**Figure 5b** lower).

#### Correlations between neural representational geometry and behaviour

To relate neural geometry to behaviour, we quantified the association between representational strength in dACC and inference accuracy. For each participant, we regressed the best group-fitted geometry and each of the three theoretical templates onto that subject’s neural RDM separately, obtained the corresponding regression coefficients (beta), and computed beta-differences. These beta-difference values were then correlated with inference-task accuracy to test whether greater similarity to the orthogonal representation predicts better behavioural performance (**Figure 5c**).

### Recurrent neural network modelling

The recurrent neural network (RNN) (**Figure 2**) was trained in a supervised learning framework to predict trial-by-trial rule switch probability for the rule inference task, implemented using the RNN frame in ‘PyTorch’ (https://www.pytorch.org) (i.e., Elman recurrent network). We employed a single hidden layer RNN with 32 hidden units, which provided a balance between sufficient representational capacity and the risk of overfitting given the limited number of trials. At each time step, the RNN produced a continuous prediction derived from its hidden state and transformed by a Sigmoid output head, thereby constraining the predictions to the probability range [0,1].

Since the goal was to simulate the representations used for human rule switching, two main inputs were given to the RNN: (i) a difficulty vector encoding the perceptual discriminability of the grating stimuli, *d*_*t*_: [1,0,0] for hard difficulty, [0,1,0] for middle difficulty, [0,0,1] for easy difficulty; (ii) a one-hot feedback vector (*f*_*t*_: positive feedback = [0, 1], negative feedback = [1, 0]) providing the reinforcement signal used to update beliefs about the covert rule. Furthermore, a subject-start flag (*S*_*t*_: 0/1) was also inputted additionally, which indicated the beginning of each subject’s data sequence (*S*_*t*_ = 1), ensuring that the recurrent state was reset at subject boundaries and preventing information leakage across individuals. Thus, the input is x̃_*t*_ = [*d*_*t*_; *f*_*t*_; *S*_*t*_]. Thus the activation ℎ_*t*_ at time *t*, takes the form of equation (2), and the previous time step ℎ_*t*-1_ is expressed as equation (3):

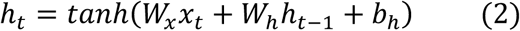

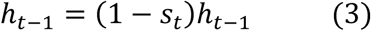

All subjects’ trial sequences were concatenated and presented jointly to the network. Hence, the RNN is trained on all the data of all subjects to obtain enough training samples. Formal trial-to-trial dependencies were maintained by the continuous propagation of the hidden state within each subject’s sequence, with resets at subject boundaries (training runs) enforced when *S*_*t*_=1. The model was trained with a batch size of 1 and optimized with the Adam optimizer (learning rate = 0.01), which adaptively scales gradient updates for stable convergence. Training was guided by a weighted mean squared error (MSE) loss function, computed at every time step. To prevent the dominance of correct trials from introducing bias into the training process, we applied a loss-weighting scheme that scaled with the number of consecutive errors. This approach encouraged the RNN to develop history-dependent evidence accumulation dynamics, thereby aligning its learning process with characteristic human behavioral adaptations and mirroring the error-monitoring functions associated with the dACC ^3^. The RNN was trained for up to 10000 epochs, with early stopping (patience = 10, delta = 0.0001) to prevent overfitting and reduce unnecessary computation. Each epoch iterated over the full dataset, allowing the network to continuously refine its mapping from difficulty and feedback history to predicted switch probability.

At evaluation, the trained network was run in ‘Pytorch eval’ mode with gradients disabled, with the same inputs and preprocessing as during training. We fed the recorded feedback from the dataset, so predictions were conditioned on the true outcome history. The activation of each unit (*h_t_*) in network hidden layer was a compact, history dependent latent code shaped by task difficulty, feedback history, and network’s internal dynamics. Thus, the unit activation can approximate the network’s internal evidence state about rule switch inference.

After training and testing the RNN, the activation of the middle layers was extracted for neural representation (patterns and geometry) analysis, and the representational patterns and geometry were visualized with the same procedure as fMRI data.

## Data and code availability

All data, code, and materials used in this study will be available at GitHub upon publication.

## Acknowledgements

This work was supported by the National Natural Science Foundation of China (Grant No. 32571283 to Q.C.) and the National Science and Technology Innovation 2030 Major Program (Grant No. 2021ZD0203800 to Q.C.). The authors thank the Brain Imaging Center at at Shenzhen University in Shenzhen, China, for assistance with MRI data acquisition. We would like to thank Ruiqi Bai, Ling Chen for their help during the data collection, and Ying Fan for helpful comments on data analysis.

## Author contributions

C.X., W.D., T.V., and Q.C. originally conceived and designed the experiments. C.X. performed the experiments. C.X., N.M., R.H., and Q.C. analyzed the data. C.X., T.V., and Q.C. wrote the paper.

## Competing interests

The authors declared have no competing interests.

**Correspondence and requests for materials** should be addressed to Qi Chen.

