## Supplementary Materials for "Orthogonal representational geometry in dACC underpins human hierarchical reasoning"

### Supplementary information for confidence-based Bayesian model

#### (1) First-stage modeling: fitting the psychometric function of perceptual judgments.

In the experiment, the test grating was physically presented with a rotation (or anti-rotation) angle of $t_s$, corresponding to the reference grating. However, the observer's internal measurement is noisy; we denote this by $\hat{t}_s$. Following hierarchical temporal-variability accounts, the observation noise is assumed Gaussian with mean 0 and standard deviation $w_m t_s$. On each trial the observer compares $\hat{t}_s$ with an internal reference threshold $\hat{t}_c$ to form $\hat{t}_d = \hat{t}_s - \hat{t}_c$ and decide clockwise vs anti- clockwise. Within a trial, $\hat{t}_c$ is treated as fixed. In addition, a "grating rotation lapse" parameter $\Gamma_t$ captures random choices of rotation direction.

Thus, the probability of pressing key "2" (denoting "right", corresponding to anti-clockwise under the red rule and clockwise under the blue rule) is:

$$p(\hat{A} = Right|\hat{C}, t_s) = \frac{\Gamma_t}{2} + (1 - \Gamma_t) \int_{\hat{t}_s} p(\hat{A} = Right|\hat{C}, \hat{t}_s, \hat{t}_c)\ p(\hat{t}_s|t_s, w_m) d\hat{t}_s \quad [1]$$

Under the red rule, if $\hat{t}_s > \hat{t}_c$ the expected "right" response has probability 1 (and 0 if $\hat{t}_s < \hat{t}_c$); under the blue rule, this mapping is reversed. The executed key press can deviate from the intended response due to "motor slip", captured by $\Gamma_{Right}$

(the probability that an intended "2" is executed as "1"). The overall probability of pressing 2 (right response) is therefore:

$$p(\mathrm{A} = Right|\hat{C}, t_s) = \frac{\Gamma_{Right}}{2} + (1 - \Gamma_{Right})\left[\frac{\Gamma_t}{2} + (1 - \Gamma_t)\int_{\hat{t}_s} p(\hat{A} = Right|\hat{C}, \hat{t}_s, \hat{t}_c)p(\hat{t}_s|\hat{t}_s, w_m)d\hat{t}_s\right] \quad [2]$$

Assuming conditional independence across trials, we obtain maximum likelihood estimates (MLE) of $\Psi_1 = \{w_m, \hat{t}_c, \Gamma_t, \Gamma_{Right}\}$:

$$\Psi_1(MLE) = \arg\max_{\Psi}\left[\sum_{i=1}^{N_{Right}} \log p(A^i = Right|\hat{C}^i, \hat{t}_s^i, \Psi) + \sum_{i=1}^{N_{left}} \log\left(1 - p(A^i = Right|\hat{C}^i, \hat{t}_s^i, \Psi)\right)\right] \quad [3]$$

Parameters were optimized with "fminsearch", and $p(\mathrm{A} = Right|\hat{C}, t_s)$ was computed as a function of rule and grating angle.

**(2) Second-stage modeling: A confidence-based model (at the rule level)**

Classical sequential-sampling models formalize choice as evidence accumulation to a decision boundary. In a hierarchical setting, causal inference about errors depends on evaluating the expected outcome given evidence quality – i.e., confidence – which humans and animals use to refine subsequent decisions. Building on this idea, we first derive an ideal-observer expression. On trial *k*, the decision to switch rules depends on the posterior odds of "switch" vs "stay", computed over the sequence of consecutive error trials $[k_0\colon k]$, where $k_0 - 1$ is the last correct trial:

$$Q_{IOM}^k = \frac{\lambda^{k_0} + \sum_{i=k_0+1}^{k} \lambda^i \prod_{j=k_0}^{i-1}(1 - \lambda^j)(1 - y^j)}{\prod_{i=k_0}^{k}(1 - \lambda^i)(1 - y^i)} \quad [4]$$

Here, $\lambda^i$ represents the subjective hazard rate (risk) of an objectively hidden rule change, and $y^j$ is the expected probability of correct perceptual discrimination on trial j (derived from the first-stage psychometric fit). The term $(1 - \lambda^j)(1 - y^j)$ represents the probability that the objective rule does not change, yet the judgment of $t_s$ was incorrect.

For simplicity and less trial number requirements, we used a confidence-based model (CBM) that approximates the ideal observer. The CBM posits a hierarchical latent variable, "switching confidence" $C_{\text{switch}}$, which characterizes the evidence for a change in the objective rule on a trial-by-trial basis. After each error trial, $C_{\text{switch}}$

increases by an amount that depends on the expected accuracy of the previous trial. A switch occurs when $C_{\text{switch}}$ crosses threshold θ. On each trial, $C_{\text{switch}}$ is assumed to be normally distributed, the mean and standard deviation denoted by $\mu_{C\text{switch}}$ and $\sigma_{C\text{switch}}$:

$$p(C_{switch}) \sim \mathcal{N}\left(\mu_{C_{switch}}, \sigma_{C_{switch}}\right) \qquad [5]$$

The CBM also assumes that the evidence for a rule switch is subject to diffuse noise. Specifically, the standard deviation of this noise increases linearly with the number of consecutive error trials and is influenced by the current trial's difficulty. By inferring $C_{\text{switch}}$ and its standard deviation, the model can thus account for how confidence, shaped by consecutive errors and difficulty, drives the decision to switch rules:

$$Q_{CBM}^{k} = \frac{P(C_{switch} \geq \theta)}{P(C_{switch} < \theta)} = \frac{\int_{\theta}^{inf} N(\mu_{C_{switch}}, \sigma_{C_{switch}}) dC_{switch}}{1 - \int_{\theta}^{inf} N(\mu_{C_{switch}}, \sigma_{C_{switch}}) dC_{switch}} = Q_{IOM}^{k} \quad [6]$$

To simplify the model so that the CBM can better fit human behavior, we assume that $C_{\text{switch}}$ =0 after any rewarded trial and θ = 1. Hence, apart from λ, the free parameters are $\sigma_{C\text{switch}}$ and α (a perseveration factor that scales $\mu_{C\text{switch}}$). The probability of switching on trial *k* becomes:

$$p(X_{y/n} = Sw|\Psi_2) = \int_{1}^{inf} \mathcal{N}(\alpha\mu_{C_{switch}}, \sigma_{C_{switch}})\, dC_{switch} \qquad [7]$$

Assuming conditional independence across trials, we obtain MLE of $\Psi_2 = \{\lambda^i, \sigma_{X_\Sigma}, \alpha\}$ by maximizing:

$$\Psi_2(MLE) = \arg\max_{\Psi} \left[ \sum_{i=1}^{N_{S_W}} \log p(X_{y/n} = S_w|\Psi) + \sum_{j=1}^{N_{NoS_W}} \log(1 - p(X_{y/n} = S_w|\Psi)) \right] \qquad [8]$$

We jointly maximized $\Psi_1(MLE) + \Psi_2(MLE)$; the AIC we report corresponds to this quantity.

### Supplementary information for psychophysical perceptual threshold task

One day before the main experiment, subjects completed the psychophysical task (Supplementary Figure 1a) for measuring subjective perceptual thresholds. In this task, a white fixation cross appeared at the center of the computer screen with a black background color, to make the subject focus on the center of screen. Then a reference grating appeared at the center of screen, followed by a short delay, and then the test grating. Subjects judged the rotation direction of the test grating compared to the reference grating, and responded by pressing the "B" key for clockwise and the "H" for anti-clockwise. Following a jittered delay range from 0.8 to 1.2 seconds, feedback was presented. The next trial started after a jittered delay ranging from 0.8 to 1.2 seconds (interval trial time, ITI). The subjects did the grating rotation judgement with 300 trials, with a rest after each 100 trials.

The logistic regression function in 'fitPsyche' toolbox [45] was used to fit the probability of subjects judging anti-clockwise of grating at each angle difference. All of our task procedure were coded with 'Psychtoolbox 3' implemented in MATALB, the 'sinGrating' function (https://bicr.atr.jp/~kmtn/imageMatlab/index.html) was used to generate reference and test gratings.

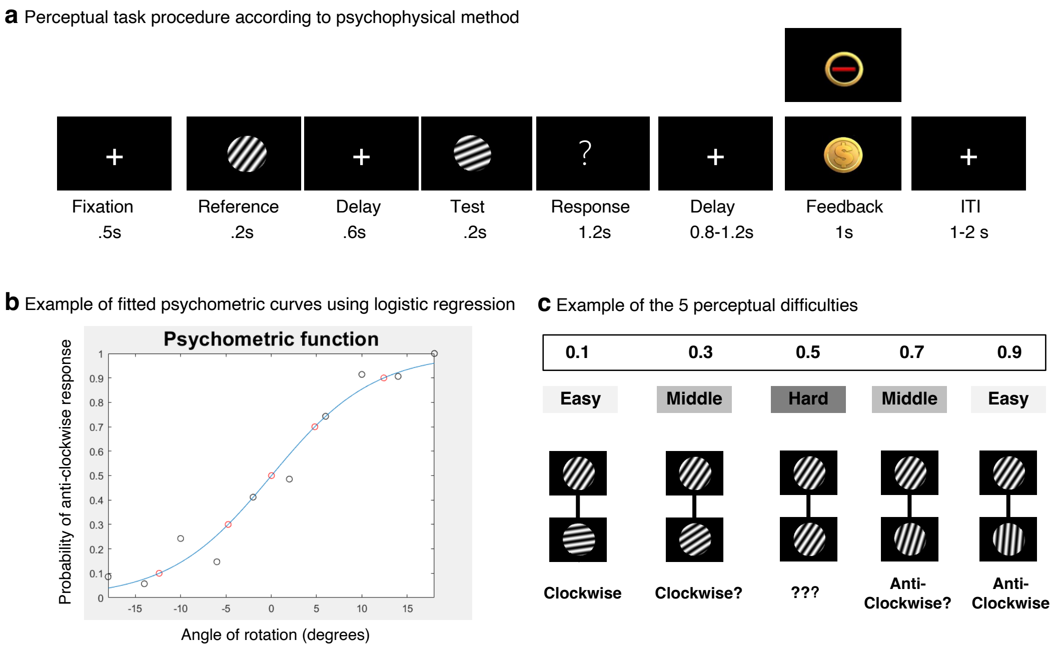


**Supplementary Figure 1. Perceptual task procedure for measuring the subjective grating rotation angle. a,** The task procedure for perceptual threshold detection. **b,** The logistic regression model fitting the psychometric curves of an example subject. Angle differences corresponding to probabilities of 0.1, 0.3, 0.5, 0.7, and 0.9 anti-clockwise judgements (red circles) were then selected as the perceptual judgement difficulty indicators in main experiments. **c,** Five probabilistic levels that subject judged as anti-clockwise, collapsed into 3 types of difficulties

(Hard, 0.5; Middle, 0.3 and 0.7; Easy, 0.1 and 0.9) was determined for each subject, and applied to the main experiment task.

### Supplementary information for computational model comparison

*Model comparison.* We compared several classical models for rule switch inference to our confidence-based model. These models focus on different aspects of rule switch inference. In the Rescorla-Wagner (RW) model, the obtained reward is used to update the value of an active stimulus-action mapping (rule) on each trial with a fixed learning rate. The adaptive learning rate (ALR) model and RW both assume that previous information is overwritten after each rule switch. The Sync model instead retains separate mappings for every task rule, and keeps track of rule switches by calculating prediction error. Model comparison confirmed that the CBM characterized subjects' behavior better than the other models.

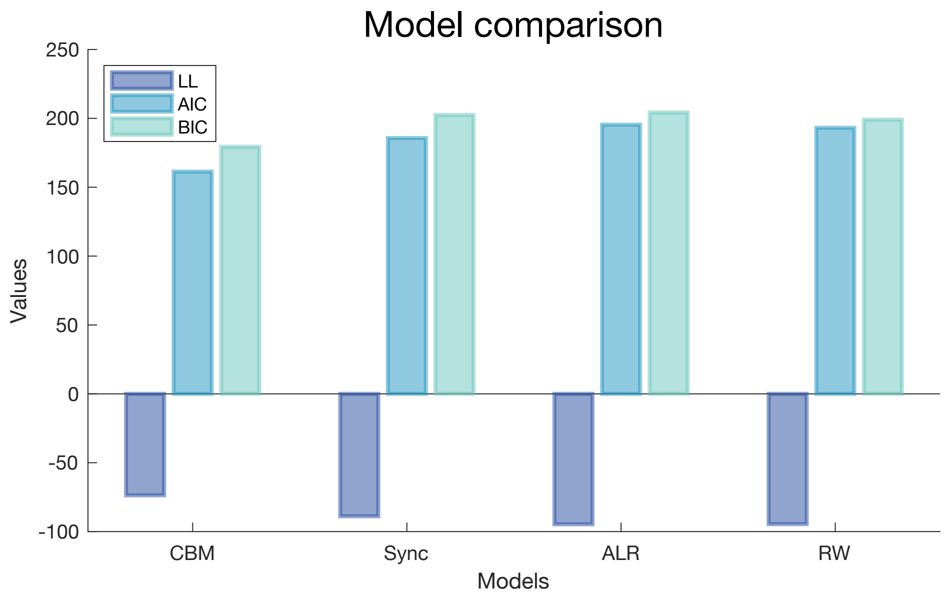


**Supplementary Figure 2. Model comparation between confidence-based model and other models of rule switch inference.** Across subjects, CBM: mean log-likelihood = -73.67, mean AIC = 161.34, mean BIC = 179.42; Sync model: mean log-likelihood = -88.88, mean AIC = 185.76, mean BIC = 202.47; ALR model: mean log-likelihood = -94.70, mean AIC = 195.41, mean BIC = 204.23; RW model: mean log-likelihood = -94.58, mean AIC = 193.16, mean BIC = 199.05. LL, log-likelihood.

### Supplementary information for univariate GLM analysis

*General Linear Model in SPM12 for task condition comparison.* We used two GLM models to detect the brain activation under different task conditions. The GLM $_{inference}$ and GLM $_{instructed}$, was fitted for rule inference and rule instructed task separately. Then two paired *t*-test was applied to detect the effects of task types (inference/instructed) and outcome type (error/reward) separately, at the group level. Multiple comparison was corrected by the cluster-based FWE (voxel level *P*<0.001) method based on random field theory in SPM12. Results of these two paired *t*-test appear in Supplementary Table 1.

**Supplementary Table 1. GLM for comparing inference and instructed rule task**

| Clusters | Anatomical Labels (AAL3) | Peak intensity | MNI coordinates | Cluster size |
|---|---|---|---|---|
| **Inference > instructed (voxel level *P* <0.001, FWE corrected cluster size 262)** | | | | |
| C1 | Insula R | 4.93 | 32 20 -4 | 317 |
| C2 | SMA_L / Frontal Sup Medial L (dACC) | 4.56 | -6 22 44 | 417 |
| C3 | Frontal Sup R/ Frontal Mid R | 4.86 | 20 26 54 | 262 |
| **Inference > instructed for error-feedback (voxel level *P* <0.001, FWE corrected cluster size 181)** | | | | |
| C1 | SMA_L / Frontal Sup Medial L (dACC) | 4.77 | -6 22 42 | 189 |
| C2 | Frontal Sup R/ Frontal Mid R | 4.87 | 20 26 52 | 181 |

*Abbreviation: SMA, supplementary motor area; dACC, dorsal anterior cingulate cortex; MNI, Montreal Neurological Institute.*

*General Linear Model in SPM12 for parameter modulation analyses.* We conducted GLMs for parameter modulation analysis of number of errors (GLM-PM1), perceptual difficulty (GLM-PM2), and confidence (GLM-PM3) separately. These parameters were modeled to modulate the feedback-evoked activation. To avoid the signal delay effects of other stimuli on feedback screen, the rule response screen (modulated by subject's rule judgement) and perceptual response screen (modulated by subject's perceptual judgement) were also modeled in the GLM-PM1-3. The statistical significance of the modulation effect was tested by a simple *t*-test against 0 at the group level, corrected for multiple comparisons by cluster-based FWE (voxel level *P*<0.001) method. GLM-PM1-3 results appear in Supplementary Table 2.

**Supplementary Table 2. GLM-PMs for modulation effect of number of errors, perceptual difficulty, confidence**

| Clusters | Anatomical Labels (AAL3) | Peak intensity | MNI coordinates | Cluster size |
|---|---|---|---|---|
| **GLM-PM1: number of errors** | | | | |
| Positive: voxel level *P* <0.001, FWE corrected cluster size 197 | | | | |
| C1 | Thalamus R | 6.85 | 10 -12 6 | 254 |
| C2 | Thalamus L | 5.82 | -14 -18 12 | 197 |
| C3 | Frontal Mid R | 4.8 | 46 34 30 | 407 |
| C4 | SMA_L / Frontal Sup Medial L (dACC) | 6.80 | -4 18 42 | 1251 |
| Negative: voxel level *P* <0.001, FWE corrected cluster size 156 | | | | |
| C1 | Amygdala R | -7.29 | 12 8 -4 | 304 |
| C2 | Hippocampus L | -5.08 | -30 -12 -18 | 328 |
| C3 | Temporal Mid L | -4.46 | -66 -16 4 | 156 |
| C4 | Cingulum Ant L / Frontal Med Orb L | -6.89 | -4 36 -4 | 2009 |
| C5 | Precuneus L/Cingulum Post L | -4.95 | -2 -46 38 | 340 |
| C6 | Angular L/Occipital Mid L | -5.64 | -42 -66 24 | 649 |
| **GLM-PM2: perceptual difficulty** | | | | |
| Negative: voxel level *P* <0.001, FWE corrected cluster size 193 | | | | |
| C1 | Frontal Inf Tri R / Frontal Mid R | -4.56 | 48 30 22 | 193 |
| C2 | SMA_L / Frontal Sup Medial R (dACC) | -4.74 | 8 20 40 | 379 |
| **GLM-PM3: confidence** | | | | |
| Positive: voxel level *P* <0.001, FWE corrected cluster size 227 | | | | |
| C1 | SMA_L / Frontal Sup Medial L (dACC) | 5.38 | -6 18 44 | 227 |
| Negative: voxel level *P* <0.001, FWE corrected cluster size 237 | | | | |
| C1 | Frontal Inf Orb L / Temporal Pole Sup L | -6.13 | -34 32 -16 | 237 |
| C2 | Temporal Sup R | -5.65 | 62 -4 -8 | 1075 |
| C3 | Temporal Mid L / Cingulum Ant l | -9.61 | -6 36 -4 | 11608 |
| C4 | Precentral R / Paracentral Lobule R | -8.37 | 8 -42 66 | 7304 |
| C5 | Angular R | -7.56 | 44 -63 27 | 729 |
| C6 | Cingulum Mid R | -5.98 | -2 10 32 | 289 |
| C7 | Frontal Sup L | -5.17 | -14 40 44 | 281 |

*Abbreviation: SMA, supplementary motor area; dACC, dorsal anterior cingulate cortex; MNI, Montreal Neurological Institute.*
